# Androgen receptor modulation of vocal circuitry in Alston’s singing mouse

**DOI:** 10.1101/2024.10.18.619097

**Authors:** Da-Jiang Zheng, Britt Mardis, Denise Lam, Tarik Islam, Joel Tripp, Steven Phelps

## Abstract

Animal courtship and aggressive displays are dramatic, often sexually dimorphic behaviors that require the coordinated modulation of diverse motivational and motor circuits. In Alston’s singing mice, a novel and elaborate advertisement vocalization is sexually dimorphic and steroid sensitive (*Scotinomys teguina*). Males sing more often than females, and on average male songs have more notes. Song is influenced by circulating androgens, but how such hormonal differences influence the diverse brain regions involved in vocal display is not understood. To characterize androgen-sensitive sites in the vocal motor pathway, we used two isoforms of pseudorabies virus (PRV) to double-label circuits ending in laryngeal and jaw muscles involved in vocalization, and co-labeled these neurons for androgen receptor (AR). Next we manipulated circulating androgens and observed the effects on AR distribution and male song. We find androgens drive coordinated changes in AR abundance across motor and motivational circuits, and both individual and group differences in song are associated with AR abundance. The results reveal how circulating androgens and the auto-regulation of androgen receptors can influence the diverse circuits necessary for elaborate advertisement displays.

**Significance statement:** Courtship and aggressive displays are among those most elaborate and dramatic of sexually dimorphic behaviors. We show that in Alston’s singing mouse, an androgen-sensitive network defined by retrograde tracing shapes vocal display effort. Our results reveal how the intensity of singing mouse vocalizations is influenced by androgen actions in the vocal motor pathway.

## Introduction

Advertisement displays are among the most dramatic and diverse behaviors observed in the animal kingdom, including the long flowing tails of the African widowbird (Andersson and Andersson 1994), the deafening choruses of American frogs (Ryan et al 1981), and the popcorn-like scent-marking of the Asian bearcat (Greene et al 2016) With their roles in courtship and aggression, advertisement displays are thought to have been driven by sexual selection, and are often sexually dimorphic. Such dimorphism relies on sex-specific patterns of morphology, motor patterns and motivation, but how such coordination is achieved is poorly understood.

Potent steroid hormones known as androgens are produced by the testes and the adrenal cortex, and are often used to coordinate the expression of vertebrate sex characters (Borg, 1994; Wang, Yeh, Tzeng, & Chang, 2009). Testosterone can be aromatized to estrogen and act through estrogen receptors, or can be metabolized to potent non-aromatizeable androgens like dihydrotestosterone (DHT) or 11-ketotestosterone, whose actions are mediated by androgen receptors. Androgens and their metabolites are involved in a broad range of sexually dimorphic behaviors, including mating, courtship and territorial displays (Schuppe et al 2022). Androgens are also important for the regulation of vocalization in many species, including songbirds (Nottebohm & Arnold, 1976), frogs (Burmeister & Wilczynski, 2001; J. Pérez & Kelley, 1996), and fish (Forlano et al., 2010) .

Vocalizations make particularly good case studies for exploring sexually dimorphic displays and their mechanisms. They are widespread and ancient among vertebrates (Chen & Wiens, 2020), and many of the neural systems that control vocalization have known homologs. Among tetrapods, limbic centers like the bed nucleus of the stria terminalis, the amygdala, and the preoptic area interact with midbrain regions like the periaqueductal gray, and lower motor nuclei to execute vocal production (Goodson & Bass, 2002; Kelley et al., 2017; Tschida et al., 2019). Perhaps surprisingly, although the vocal circuitry of mammals is well studied (Uwe Jürgens, 2002), and mammalian vocalizations are often sexually dimorphic (Charlton & Reby, 2016), little work has examined the role of androgens on mammalian vocal circuits or its relationship to dimorphic vocalization. Here we ask how androgens act upon the vocal apparatus and circuitry to shape dimorphic vocal behavior in Alston’s singing mouse.

Alston’s singing mouse (*Scotinomys teguina*) is a small muroid rodent in the Cricetidae family that vocalizes in a frequency range audible to humans (Campbell et al., 2010; Hooper & Carleton, 1976; Miller & Engstrom, 2007). The advertisement song of *S. teguina* consists of a rapid series of downward frequency sweeps that get progressively longer over the ∼7s of a song (Burkhard et al., 2018; Campbell et al., 2010; Giglio & Phelps, 2020). The song is used for female attraction as well as male-male competition (M. Fernández-Vargas et al., 2011; Pasch, George, Campbell, et al., 2011b). Singing is sexually dimorphic: males sing more often than females, and their songs tend to last longer (Miller & Engstrom, 2007, Tripp and Phelps 2024). Castration of males makes them sing less often, and with songs that are shorter in duration – an effect that can be rescued by dihydrotestosterone (DHT; Pasch, George, Hamlin, et al., 2011). Recently, we used two isoforms of pseudorabies virus, a transsynaptic retrograde virus, to map the neural inputs to the larynx and jaw of these animals (Zheng et al 2022). We found broad overlap of the vocal circuits with regions that exhibit androgen-receptor immunoreactivity.

Here we triple label for androgen receptor and for PRV isoforms injected into the larynx and jaw to assess whether the specific neurons that participate in the vocal circuit are androgen sensitive. Next, we perform a castration/replacement study to examine the effects of androgens on singing behavior. Finally, we collected brains and quantified AR expression in a series of regions related to vocal behavior and motivation. Together these data allow us to examine the relationship between dimorphic aspects of song and the actions of androgens on vocal networks.

## Methods

### Animals

For our two experiments, we used a total of 19 male *S.teguina* outbred from a wild population caught near Quetzal Education Research Center (QERC), in San Gerardo de Dota, Costa Rica. Subjects were fed an *ad libitum* mixture LabDiet Feline and LabDiet Insectivore chow (LabDiet, St Louis, MO, U.S.A.). Animals were housed with same-sex sibs at 12h light:dark cycles and substantial enrichment (Zheng et al, 2021). All subjects were sexually mature and gonadally active at the time of the experiments. *S.teguina* weighed 13.0±1.67 grams and were 94.7±12.3 days of age at the time of gonadectomy and hormone implantation. Animal protocols were approved by the IACUC committee at The University of Texas at Austin in accordance with the National Institute of Health *Guide for the Care and Use of Laboratory Animals*.

### Identification of androgen-responsive vocal circuits

As previously described (Zheng et al, 2022), we employed two isogenic trans-neuronal retrograde viral tracers, one with a GFP reporter and the other with an RFP reporter. One isoform targeted the cricothyroid muscle of the larynx, a muscle essential to frequency modulation of vocal behavior (Smith 2024), and the other targeted the anterior digastricus of the jaw, which modulates the opening of the mouth with each note (Okobi et al 2019). 96 hours after inoculation with both PRV viral constructs (n=5), we perfused each subject, dissected out the brain, cryoprotected through sucrose, and then froze the brain and stored it at −80°C until sectioning.

### Hormone manipulation

Hormone implants were constructed similar to our previous report (Pasch, George, Hamlin, et al., 2011). Briefly, male mice were anesthetized with isoflurane and castrated bilaterally through two paramedian incisions flanking the scrotal region. Incisions were then glued using Gluture (Zoetis). We then placed a 10-mm silastic implant along the dorsal midline. Implants were filled with either 1mm of cholesterol (Sigma C8667, n=8) or dihydrotestosterone (DHT, Cayman 15874, n=6). (Cholesterol is a precursor for steroidal hormones that serves here as a control substance (Wilson 1996)) Implants were sealed with silicon adhesives and disinfected with chlorohexidine before insertion into subject animal. Animals were then isolated in a clean cage and observed every 12 hours for the first two days and then daily for the first week.

### Behavioral testing

Thirteen days post-surgery, animals were introduced to a playback chamber (Giglio & Phelps, 2020). Animals were placed in 61 × 45.7 cm and 25.4 cm tall plexiglass containers that contained the same food, water and enrichment items as their home cages. Animals were singly housed in acoustic-attenuating chambers 40 hours before playbacks began. Subjects were recorded throughout the 40 hours using equipment previously reported (Giglio & Phelps, 2020) with a custom MATLAB code that allowed for continuous recording of the subjects.

All playbacks occurred between 10-12h in the morning. Thirty minutes before playbacks began, all enrichment except water and food were removed. The playback procedure has been described in Giglio & Phelps (2020). Briefly, a master timer file was created with 20 timeslots randomly distributed within an hour. For each timeslot, one of four different songs recorded from a single male was broadcast to the subject. Songs were selected from a library of *S.teguina* songs recorded from the males collected in the wild (Burkhard et al., 2018), and were from animals unfamiliar to the experimental subjects. Songs used as stimuli in this experiment represented approximately median song duration (Burkhard et al., 2018).

### Perfusion and tissue dissection

Animals were euthanized with isoflurane, to effect, and then subjects were transcardially perfused with a peristaltic pump (Cole-Parmer), first with cold 1X PBS (Invitrogen) and then cold 4% paraformaldehyde (EMS) in 1X PBS (Zheng et al, *2021*). Brains were dissected out, post-fixed in 4% paraformaldehyde for 24 hours in a cold room and then cryoprotected in 15% and then 30% sucrose. For our PRV study, we perfused at 96-hours post-inoculation. For our behavioral study, perfusion began one hour after the playback window. After cryoprotection, brains were frozen on powdered dry ice and stored in −80°C freezer until sectioning.

### Immunofluorescence

For the PRV study, fixed and frozen brains were sectioned onto slides and triple-labeled for GFP, RFP, and AR. We sectioned 30um alternative sections onto two series of plus charged slides. One series went into triple labeling experiment with anti-GFP antibody (1:1000), anti-RFP antibody (1:500) and PG-21 (1:1000). Following Zheng et al (2022), we washed sections and blocked with normal serum. Sections were then incubated with primary antibodies for 24 hours in room temperature. They then were rinsed again and then incubated in secondary for 2 hours. Slides were then covered with mounting media.

Based on previous experiments (Zheng et al, *2021*, *2022*) we narrowed our focus to a qualitative assessment of the presence of triple-labeled cells within five brain regions implicated in vocalization. The bed nucleus of the stria terminalis (BST) and medial preoptic area (MPOA) were selected based on their roles in diverse motivated social behaviors, including mating and courtship (Newman 1999, Crews and Moore 1985, Goodson 2008, O’Connell and Hofmann 2012). The lateral periaqueductal grey is known to gate mammalian vocalization, while the nucleus ambiguous contains motor neurons that drive laryngeal muscles (Hisa et al 1984). Lastly, the gigantocellular reticular formation was identified as a putative central pattern generator for the singing mouse vocalization, but does not express androgen receptor (Zheng et al. 2021, 2022). To examine the AR labeling, we selected the 640nm channel for focus. For 20X visualization we used a Plan Apochromat λ 20X, NA 0.75 objective, for 40X we used a Super Plan Fluor ELWD 40X; NA 0.6 objective. Otherwise, microscopy is done as previously described (Zheng et al 2022)

For the behavioral study, frozen brains were first embedded in OCT mounting medium and then sectioned at 30μm thickness from the appearance of the secondary motor cortex to the nucleus ambiguous. Sections were mounted on Superfrost Plus slides (Fisher Scientific) and placed at −80°C until labeling. For immunofluorescence of the androgen receptor, sections were taken out of the freezer and allowed to dry before lining the border with a hydrophobic barrier pen (Abcam). Sections were then incubated with PG-21 at 1:1000 (EMD Millipore) for 24 hours and then washed. A goat anti-rabbit alexa fluor 594 secondary (Thermofisher) was used to bind to the primary. After another set of washes, slides were mounted on coverslips with Prolong Diamond with DAPI (Thermofisher).

Microscopy of both DAPI and AF594 was done using the same apparatus as above (Zheng et al 2022) with images taken at 20X for the five ROIs. Laser intensity and gain were kept consistent throughout the microscopy session. One ROI was taken for all subjects continuously within one session.

### Acoustic analysis

Acoustic analysis followed prior studies (Burkhard et al., 2018; Campbell et al., 2010; Giglio & Phelps, 2020). Briefly, all songs were analyzed by custom code in MATLAB (Campbell et al., 2010). The songs of *S.teguina* consists of downward frequency sweeps (notes) that increase in number, amplitude, and bandwidth as the song progresses. The code first identifies the boundaries of notes by identifying amplitudes three standard deviations above background. The start and stops of notes are used to measure note durations and internote intervals. Each individual note is then analyzed spectrally using an FFT, and the descending frequency sweep of each note is described by a quadratic curve with three paramaters – where FMc denotes the intercept, or starting frequency of the note, FMb denotes the starting slope of the note, and FMa denotes the note curvature. The frequency and amplitude modulation of a given note, including its duration and amplitude envelope, change systematically as the song progresses, and so each note feature can be fit to a quadratic formula that describes the note as a function of its position in the song. The starting frequency of the note is thus described by the song-level quadratic defined by parameters FMc_C, FMc_B, and FMc_A; the note slope by FMb_C, FMb_B, and FMb_A; and the note curvature by FMa_C, FMa_B, and FMa_A. Lastly, we calculate a variety of features from entire songs. For spectral characteristics, these include the minimum frequency (MinHz), the maximum frequency (MaxHz), and bandwidth (BW = MaxHz - MinHz), each calculated from the fundamental frequencies of notes. For amplitude characteristics, global song measures include mean song length (seconds), mean note number, and lastly, we examined the total number of songs in a playback hour.

### Cell counting

Cell counts were quantified using CellProfiler 4.2.5, a cell image analysis software. Images were first adjusted in FIJI using a custom macro (split_adjust_merge.ijm). The DAPI channel was automatically adjusted for brightness and contrast to maximize visibility of the cells. The red channel was then adjusted to the same brightness and contrast parameters, functionally normalizing their brightness to that of the DAPI label. The two channels were then merged used in the analysis pipeline. Pipeline inputs consisted of merged color images produced as above. The pipeline created an illumination function averaged from all the merged input images to correct for illumination differences across the field of view. The original color image was then split back into separate red, and blue channels, which were each converted to grayscale. The illumination function was then applied to each channel individually. Cells were identified using an adaptive Robust Background thresholding method. This calculates a different pixel intensity threshold for each pixel, thus adapting to changes in foreground/background intensities across an individual image. For each pixel, the threshold is calculated based on the pixels within a given neighborhood (or window) surrounding that pixel. After an initial pipeline run containing all the images for a given region, images were manually inspected. Images were flagged for re-processing based on the presence of debris, ventricles, interfering fiber tracts, or poor performance by the thresholding algorithm. These flagged images, as well as the ones flagged automatically for oversize cells, were then re-processed one region at a time using the same parameters as before, except with the addition of a module that allowed the experimenter to manually edit identified objects, removing erroneous identifications and adding additional objects where necessary. This editing process was done blind to experimental treatment and subject identity. All images for a single region (across individual mice) were processed together as a batch.

### Statistical Analyses

All analyses were performed using R (v4.2.2). Comparisons of song measures (counts, mean duration, and mean note number) and AR+ cell counts across treatment groups were made using two-sided t-tests. Animals that sang zero songs during the playback period were excluded from comparisons of mean duration and mean note number. Pearson’s correlations were used to compare AR+ cell counts with song measures.

## Results

### AR immunoreactivity within song circuits

In the lateral periaqueductal grey (LPAG; Figure 1b1-1b3), medial bed nucleus of the stria terminalis (BNSTm; Figure 1c1-1c3) and the medial preoptic area (MPOA, Figure 1d1-1d3), we found multiple cells within each region that were positively labeled for all three markers (Larynx PRV, Jaw PRV, and AR+). In laryngeal motor neurons of the nucleus ambiguus (AmB; Figure 1a1-1a3), we found cells were labelled with two markers (Larynx PRV and AR) but were not labelled with PRV injected into the jaw. Neurons in the gigantocellular reticular formation (Gi) exhibited double labelling for both jaw and laryngeal PRV markers, but we did not find concentrated punctate nuclear androgen receptor expression.

**Figure 1:**
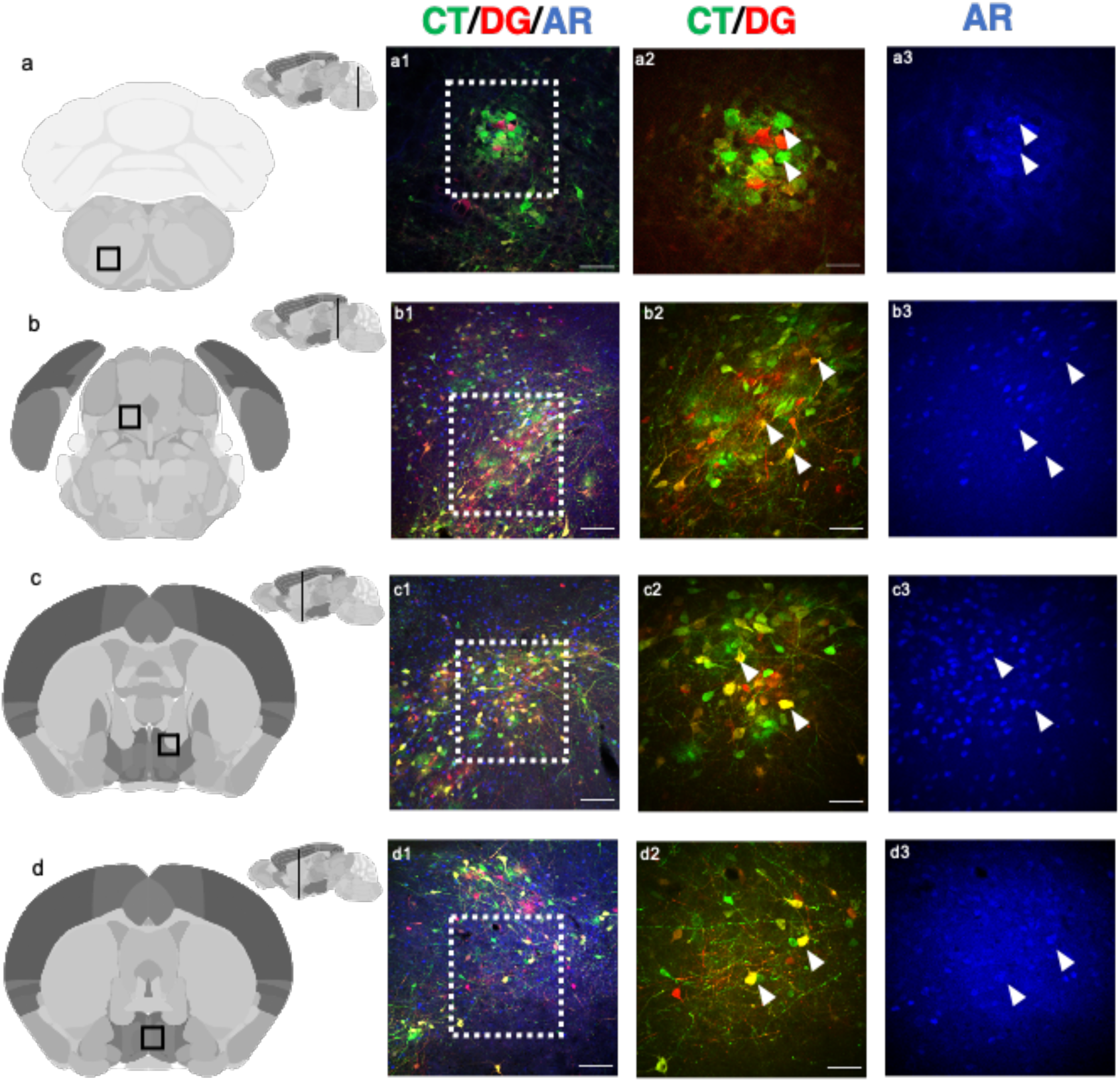
Areas of triple-labeled (co-labeled for PRV, pseudorabies virus, and AR, androgen receptor) in *S.teguina* brain. Figure 1a, nucleus ambiguus (AmB). Figure 1b, lateral periaqueductal grey (LPAG). Figure 1c, medial bed nucleus of the stria terminalis (BNSTm). Figure 1d, medial preoptic area (MPOA). Scale bars for “CT/DG/AR”= 100μm. Scale bars for “CT/DG” = 50μm (CT, injection into the cricothyroid/larynx; DG, injection into the digastricus/jaw; AR, androgen receptor)

### Song rates

DHT implanted individuals sang more songs (Figure 2a; Welch’s, Cholesterol median= 4.5; DHT median= 12.83; t=-4.1; p=0.0021) that had more notes (Figure 2b; Welch’s, Cholesterol median= 73; DHT median= 88; t=-2.5; p=0.038) and were longer in duration (Figure 2c; Welch’s, Cholesterol median=5.8; DHT median= 7.6; t=-2.7; p=0.020) in the playback hour with sample sizes of Cholesterol n=7 and DHT n=6 (one cholesterol individual did not sing within the playback hour). Overall song duration was strongly predictive of note number (r=0.92).

**Figure 2.**
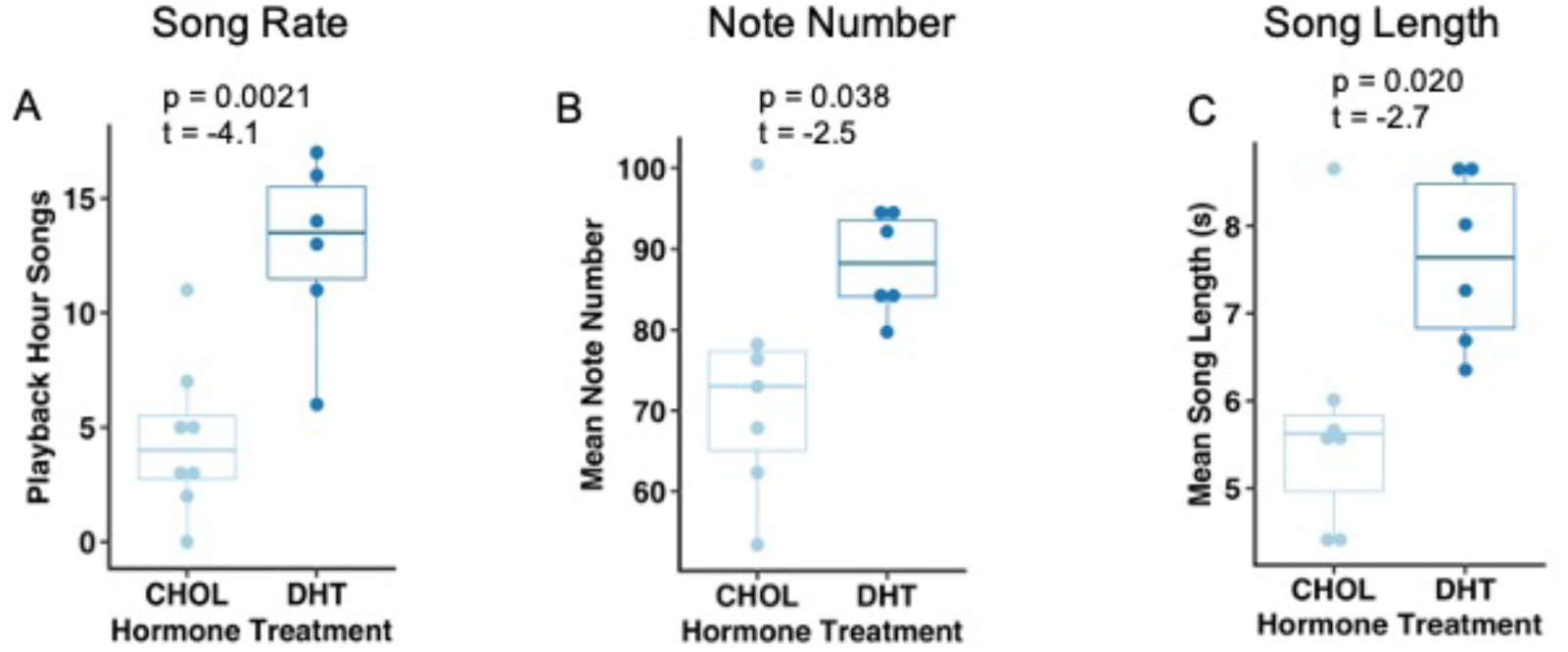
Song effort comparisons between implanted groups. Box and whisker plot for song effort measurements between cholesterol treated animals (n=8) and DHT treated animals (n=6). Figure 2a, number of songs in the playback hour. Figure 2b, mean number of notes per song. Figure 2c, mean length of songs, in seconds, within the playback hour. CHOL, cholesterol treated animals. DHT, dihydrotestosterone treated animals.

### Song frequency modulation

We measured twelve spectral properties for the songs, including maximum frequency, minimum frequency, mean bandwidth, FMa_a, FMa_b, FMa_c, FMb_a, FMb_b, FMb_c, FMc_a, FMc_b, and FMc_c. None of the 12 spectral properties were significantly different between the two groups. (Table 1).

**Table 1.** Frequency modulation and spectral characteristics. Comparison between cholesterol and DHT implanted animals a subset of frequency modulation traits and spectral properties. All trait comparisons here are non-significant between the two groups.

| Song parameter | T value | P value |
| --- | --- | --- |
| Mean Bandwidth | -1.14 | 0.28 |
| Min frequency | -1.15 | 0.27 |
| Max frequency | 2.01 | 0.07 |
| FMa_a | 0.34 | 0.74 |
| FMa_b | 0.07 | 0.95 |
| FMa_c | -0.48 | 0.64 |
| FMb_a | -1.65 | 0.13 |
| FMb_b | 1.24 | 0.24 |
| FMb_c | -0.45 | 0.66 |
| FMc_a | 0.86 | 0.41 |
| FMc_b | -0.77 | 0.46 |
| FMc_c | 1.49 | 0.17 |

### Androgen receptor group differences

There were strong DHT vs CHOL differences in AR abundance in four of the brain regions that we analyzed, including the nucleus ambiguus (t=-14.60, p=2.9e-06),lateral periaqueductal grey (t=-3.5, p=0.011), bed nucleus of the stria terminalis (t=-5.7, p=0.0011), and the medial preoptic area (t=-4.44, p=0.0050). As mentioned, the reticular formation did not contain androgen receptor immunoreactivity, and AR cell abundance between the two treatment groups was not significantly different (t=-1.3, p=0.25) in this region.

### Androgen receptor abundance predicts song effort

Sex differences in song effort are reflected in both the number of songs sang in an hour (song rate), and in the duration of individual songs (measured in song duration (s), or note number). AR abundance was strongly predictive of song rate in all four of the AR/PRV+ regions we analyzed, including the nucleus ambiguus (r=0.81, p=0.00050, Figure 3a), lateral periaqueductal grey (r=0.57, p=0.035, Figure 3b), medial bed nucleus of the stria terminalis (r=0.73, p=0.0030, Figure 3c), and the medial preoptic area (r=0.68, p=0.0075, Figure 3d). AR+ cell counts in three of the four regions – nucleus ambiguous, MPOA and mBST – were significantly correlated with song duration (Table 2). Neither song rate nor song duration were significantly correlated with AR abundance in the reticular formation (Figure 3e).

**Figure 3.**
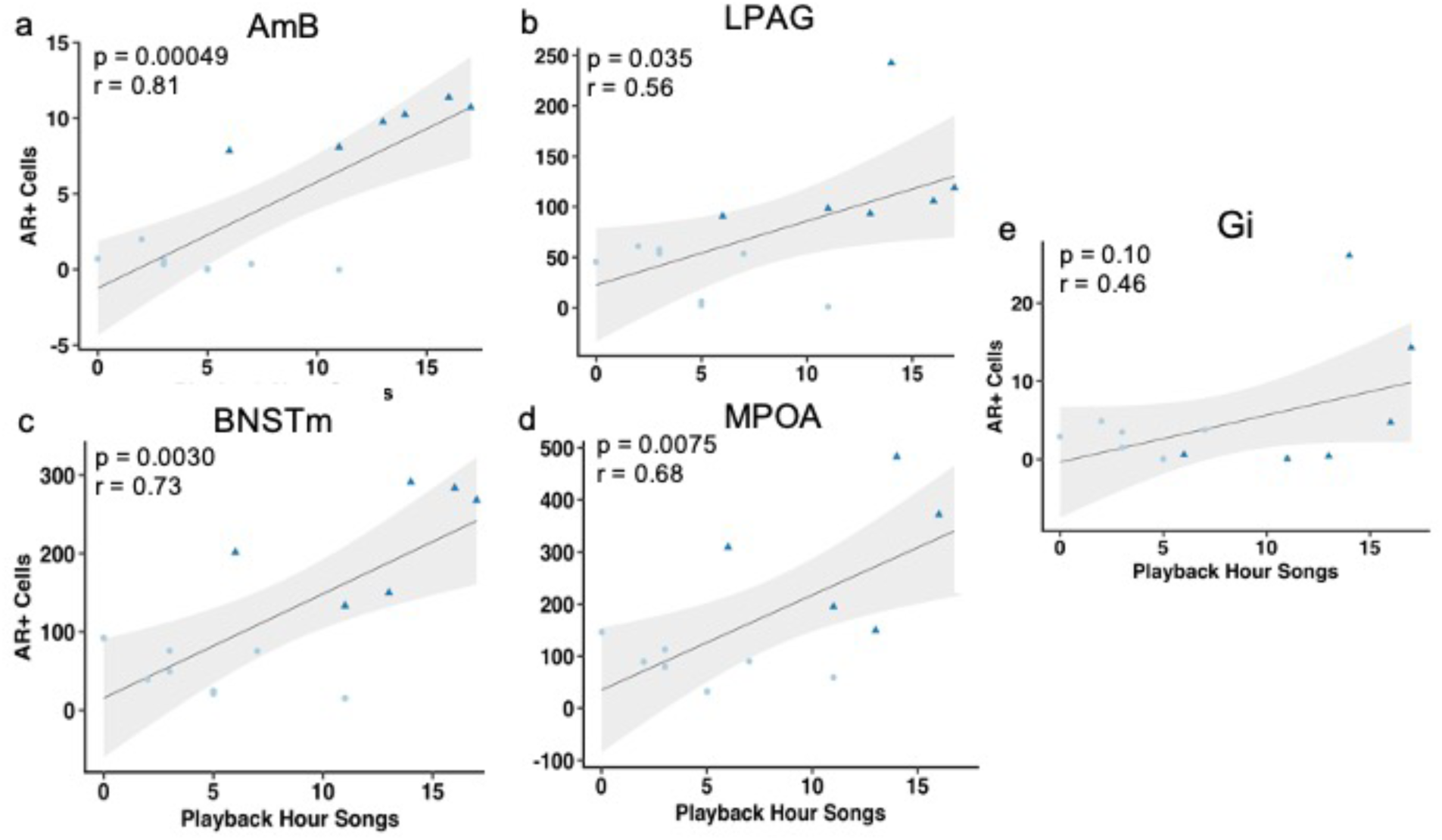
AR+ cells in various ROIs with relationship to song rate. Pearson’s correlation graph of all ROIs with song rate within the playback hour. Figure 3a, nucleus ambiguus. Figure 3b, lateral periaqueductal grey. Figure 3c, medial bed nucleus of the stria terminalis. Figure 3d, medial preoptic area. Figure 3e, gigantocellular reticular formation.

**Table 2.** AR+ cells in various ROIs with relationship to other measures of song effort. Pearson’s correlation of r and p value are reported for each ROI. AR, androgen receptor; ROI, region of interest; AmB, nucleus ambiguus; LPAG, lateral periaqueductal grey; BNSTm, medial bed nucleus of the stria terminalis; MPOA, medial preoptic area; Gi, gigantocellular reticular formation.

| AR+ Cells in ROI | Mean Note Number |  | Mean Song Length |  |
| --- | --- | --- | --- | --- |
|  | r value | p value | r value | p value |
| AmB | 0.56 | 0.046 | 0.61 | 0.027 |
| LPAG | 0.45 | 0.12 | 0.37 | 0.21 |
| BNSTm | 0.53 | 0.06 | 0.77 | 0.0022 |
| MPOA | 0.59 | 0.034 | 0.64 | 0.018 |
| Gi | 0.38 | 0.20 | 0.27 | 0.38 |

## Discussion

In the current study, we characterized androgen receptor distribution in neurons of the vocal motor circuit, and manipulated androgens to examine how this sex steroid might contribute to sex differences in the vocal behavior of singing mice. We previously identified regions that reside within the vocal circuit based on PRV tract-tracing, and found that these same regions are often positive for androgen receptor immunoreactivity (Zheng et al 2022). Here we find that in regions with PRV+ and AR+ neurons, these labels commonly co-localized to the same neurons. To examine the effects of androgens on singing behavior, we performed a castration and androgen-replacement experiment, which revealed that androgens promote sexually dimorphic aspects of *Scotinomys* song. Dihydrotestosterone had a pronounced effect on measures of song effort and motivation, including song rate and length. Similarly, we find that AR abundance across brain regions is also strongly predictive of individual differences song rate and duration. Adult androgen manipulations thus recapitulate patterns of sexual dimorphism in song. Notably, we found no effects of androgens or AR staining on patterns of frequency modulation, aspects of song that differ between individual mice, but that are not known to be sexually dimorphic. We now examine these findings in more detail.

In a prior study, we used retrograde double-labelling of neurons with PRV isoforms targeted to jaw and laryngeal muscles to elucidate a vocal circuit in singing mice (Zheng et al. 2022). Here, we examined four regions that were AR+ and exhibited labeling for PRV. These include the bed nucleus of the stria terminalis (BNST), medial preoptic area (MPOA), lateral periaqueductal grey (LPAG), nucleus ambiguus (AmB). Both the BNST and MPOA shape motivation (Berridge, 1996) particularly with respect to socio-sexual behavior and aggression (Albert, Walsh, Gorzalka, Siemens, & Louie, 1986; Edwards, Nahai, & Wright, 1993; Fuxjager, Forbes-lorman, Coss, Auger, & Auger, 2010; Liu, Salamone, & Sachs, 1997). At the level of the midbrain, the LPAG is thought to gate a specific set of vocal behaviors in mammals, including lab mice (Gruber-Dujardin, 2010; Tschida et al., 2019). This region also shows an abundance of nuclear labeling. We also note that the presence of androgen receptors in the nucleus ambiguus and PRV+ for the larynx infection are concentrated near the “cricothyroid compartment” (Hisa et al., 1984). Overall, our anatomical work suggests that for a broad range of regions within the vocal circuit that we examined, PRV and AR immunoreactivity are co-occurring within the same neurons. The candidate central pattern generator (CPG) from our previous work is the gigantocellular reticular nucleus, and it stands out as an exception to the overall pattern we report. It does not show clear nuclear staining for androgen receptors.

We find that a non-aromatizeable androgen, DHT, promotes “song effort” traits of the *S.teguina* vocalization (Burkhard et al., 2018; Giglio & Phelps, 2020; Pasch, George, Hamlin, et al., 2011). Specifically, DHT subjects increased song rate as well as song length as measured either in seconds or in number of notes. These findings mirror previous observations of sexual dimorphism in song rate and length of the *S.teguina* vocalization reported previously (Campbell et al., 2010; Miller & Engstrom, 2007; Tripp and Phelps 2024) as well as effects of androgens on “song performance” of this species (Pasch, George, Campbell, et al., 2011a). Broadly speaking, our study along with others support that androgens drive sexually dimorphic aspects of singing behavior (Fischer & Kelley, 1991; Gahr & Metzdorf, 1999; Pasch, George, Hamlin, et al., 2011).

In general, finding that androgens regulate song effort in singing mice is consistent with a more general perspective that androgens signal reproductive context, tying life-history stage to increased rates of display (Monks & Holmes, 2018; Tobiansky & Fuxjager, 2020). Similar findings have been reported in barn owls (Béziers, Ducrest, Simon, & Roulin, 2017), canaries (Alward, Balthazart, & Ball, 2017), golden-collared manakins (Fuxjager, Miles, Goller, Petersen, & Yancey, 2017), rock frogs (Mangiamele et al., 2016) and cichlid fish (Alward et al., 2019). From a mechanistic perspective, in many taxa sex differences in motivation are reflected in AR distribution in the forebrain and midbrain. The BNST, MPOA, and LPAG are critical for sociosexual motivation and are androgen-dependent across vertebrates (Goodson, 2005; O’Connell & Hofmann, 2011). Similarly, the MPOA is generally important for courtship and display across vertebrates (Alger & Riters, 2006; Jacques Balthazart & Ball, 2007; Wade, Huang, & Crewst, 1993). More specifically, the MPOA and LPAG are both known important for vocalizations of lab mice (Gao, Wei, Wang, & Xu, 2019; Tschida et al., 2019, Michael and Tschida 2020). Since these same structures seem to be part of an ancient vocal circuit, the data suggest that these regions are an ancient source of sexual dimorphism in animal displays.

In singing mice, we find that DHT had no apparent influences on spectral properties of song. We showed no significant differences between the two groups in max HZ, min HZ, and bandwidth. The mechanisms of sound production of singing mice and their close relatives the pygmy mice are thought to involve a laryngeal whistle, with frequency components shaped by the size of the ventral pouch and the contraction of the laryngeal cricothyroid muscle (Riede & Pasch, 2020, Smith et al 2021), one of the targets of our PRV tract tracing. A limited study suggests no sex differences in the structure of the larynx (Smith et al 2021), and is consistent with our findings.

The songs of singing mice are highly frequency modulated (Campbell et al., 2011). Measures of frequency modulation that we describe as “note shape” change systematically over the song as notes elongate, and are individually distinctive (Burkhard et al., 2018; Giglio & Phelps, 2020). Despite abundant and repeatable individual differences, there do not seem to be sex differences (TT Burkhard, *personal communication*). Similarly, we found no effects of castration or DHT replacement on these aspects of note shape. This is consistent with the putative role of note shape in identity signaling (Burkhard et al., 2018). We assume note shape, which requires the coordination of a variety of muscles in precise order – muscles such as the cricothyroid, jaw, intercostal and diaphragm muscles – is managed by a CPG. It is remarkable that the CPG suggested by our PRV work, the gigantocellularis (Gi) stands out as relatively unique within the vocal circuit as lacking androgen receptors. Thus, this negative finding is consistent not only with the lack of sexual dimorphism in frequency modulation, but in the distribution of androgen receptors in the vocal circuit overall. It is not clear why this should be an exception, particularly since the identification of sex would seem to be useful for vocalizing animals. One possible explanation is that the CPG governing song is coupled to respiratory rhythms, as evident, for example, when singing mice take a shallow breath with each note (Pasch et al 2011, Okobi et al 2019). Such respiratory coupling may constrain the ability of the CPG to exhibit sex specific function.

Interestingly, *Xenopus* frogs vocalization is controlled by an androgen-sensitive CPG that is thought to be newly derived independent of respiratory muscles, while a distinct and more ancient respiratory CPG shared with females (Zornik and Yamaguchi 2008). The definitive characterization of the *Scotinomys* vocal CPG will be essential to testing this and alternative hypotheses.

All the regions of interest expected to be involved in song motivation and gating (AmB, LPAG, MPOA, and BNST) showed a significant relationship between AR expression and song rate during the playback period. One mechanism that can explain this relationship is autoregulation of the AR protein itself, in which the binding of androgens to their receptors upregulate the expression of the androgen receptor gene (Lu et al 1998, Fuxjager et al 2010). AR autoregulation is known in both the MPOA and BNST (Handa et al 1986, Lu et al 1998), but our experiment suggests such autoregulation throughout AR+ brain regions.

Overall our data suggests that androgens are acting upon multiple nodes within *S.teguina* vocal circuits to influence neural activity related to song effort. These results reveal how individual differences in behavior can emerge from the coordinated modulation of neural circuits. Together our work will help understand the integrated evolution of complex and sexually dimorphic behaviors.

## Notes

### Competing Interest Statement

The authors have declared no competing interest.

